# Capsular polysaccharides cross-regulation modulates *Bacteroides thetaiotaomicron* biofilm formation

**DOI:** 10.1101/2020.03.25.005728

**Authors:** Nathalie Bechon, Jovana Mihajlovic, Sol Vendrell-Fernández, Florian Chain, Philippe Langella, Christophe Beloin, Jean-Marc Ghigo

## Abstract

*Bacteroides thetaiotaomicron* is one of the most abundant gut symbiont species, whose contribution to host health through its ability to degrade diet polysaccharides and mature the immune system is under untense scrutiny. By contrast, adhesion and biofilm formation, which are potentially involved in gut colonization, microbiota structure and stability, have hardly been investigated in this intestinal bacterium. To uncover *B. thetaiotaomicron* biofilm-related functions, we performed a transposon mutagenesis in the poor biofilm-forming reference strain VPI 5482 and showed that capsule 4, one of the eight *B. thetaiotaomicron* capsules, hinders biofilm formation. We then showed that the production of capsules 1, 2, 3, 5 and 6 also inhibits biofilm formation and that decreased capsulation of the population correlated with increased biofilm formation, suggesting that capsules could be masking adhesive surface structures. We also showed that, by contrast, capsule 8 displayed intrinsic adhesive properties. Finally, we demonstrated that *BT2934*, the *wzx* homolog of *B. thetaiotaomicron* glycosylation locus, competes with capsule production and therefore impacts its adhesion capacity. This study identified regulation of capsular polysaccharides as a major determinant of *B. thetaiotaomicron* biofilm formation, providing new insights into how modulation of different *B. thetaiotaomicron* surface structures affect *in vitro* biofilm formation.

## INTRODUCTION

*Bacteroides thetaiotaomicron* is an abundant bacterial symbiont of the normal mammalian intestine that contributes to shaping the nutrient environment of the gut microbiome through degradation of complex polysaccharides and production of short chain fatty acids (1–5). *B. thetaiotaomicron* was also shown to stimulate the development of gut immunity (6), attenuate intestinal inflammation (7) and to strengthen the intestinal protective barrier (8, 9). Consistently, decrease in abundance of *B. thetaiotaomicron* and other *Bacteroides* species has been correlated with gut inflammation and disease emergence, underlining the importance of the gut microbiota for host intestinal physiology and health (10). By contrast, microbial functions involved in the establishment and maintenance of a healthy gut microbiota are still not well understood. It is speculated that the ability of symbiont bacteria to form biofilms could contribute to microbiota stability (11, 12). However, although bacterial biofilm formation has been studied in various facultative symbiotic and pathogenic anaerobes, information on this widespread lifestyle is still scarce in *B. thetaiotaomicron* (13–15). Whereas a comparative gene expression profiling between biofilm and planktonically grown *B. thetaiotaomicron* showed biofilm-associated up-regulation of polysaccharide utilization systems and capsule 8, one of the eight *B. thetaiotaomicron* capsule synthesis loci (15, 16) there is still no direct proof of the contribution of these surface structures to adhesion and biofilm formation. We recently showed that, although biofilm capacity is widespread among *B. thetaiotaomicron* isolates, the widely used reference strain VPI 5482 is a poor biofilm former. Nevertheless, use of a transposon mutagenesis followed by a positive selection procedure revealed mutants with significantly improved biofilm capacity, due to alteration of the structure of a putative type V pilus (13). In this study, we showed that regulation of capsule expression is another major determinant of biofilm formation by masking or unmasking adhesive *B. thetaiotaomicron* structures. This study provides new insights into the roles of capsular polysaccharides in *B. thetaiotaomicron* and their impact on the physiology and biofilm formation of a prominent gut symbiont.

## RESULTS

### Transposon insertion in capsule 4 biosynthesis operon promotes *B. thetaiotaomicron* biofilm formation

Among the previously identified transposon mutants displaying increased *in vitro* biofilm formation capacity compared to the wildtype *B. thetaiotaomicron* VPI5482 (WT) (13), 5 of them corresponded to insertions within capsule 4 (*CPS4*) synthesis operon *BT1358-1338*, encoding one of the eight capsular polysaccharides of *B. thetaiotaomicron* (Figure 1AB and table S1) (16, 17). To confirm the increased biofilm phenotype of the transposon mutants, we deleted all 19 *CPS4* structural genes located downstream of the regulators *BT1358-1357*. Crystal violet staining of *in vitro* biofilm formed in 96-well microtiter plates showed that the resulting *ΔBT1356-1338* mutant (hereafter named *ΔCPS4*) displayed a significant increase in biofilm formation compared to the wild type *B. thetaiotaomicron* VPI 5482 (Figure 1C).

**Figure 1.**
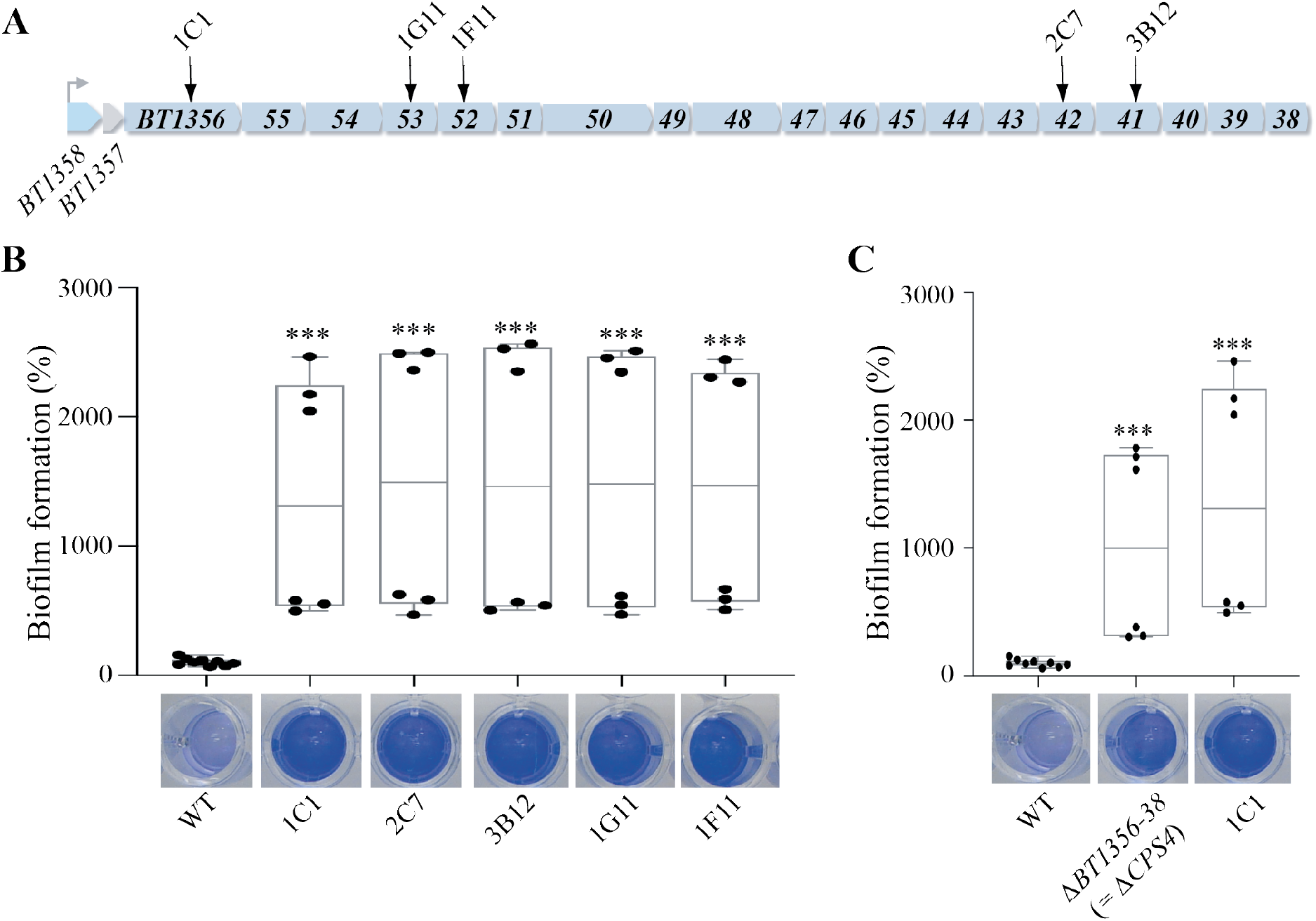
Capsule 4 inhibits biofilm formation in *B. thetaiotaomicron* VPI5482. **A.** Organization of *B. thetaiotaomicron* capsular operon 4 (*CPS4*). The first two genes (*BT1358* and *BT1357*) code for regulators of capsular biosynthesis. *BT1356-1338* code for the enzymes involved in Cps4 capsular polysaccharide biosynthesis. Arrows indicate 5 individual transposon insertions within the *CPS4* operon. **B.** 96-well plate biofilm assay after 48h growth in BHIS. Mean of WT is adjusted to 100 %. Min-max boxplot of 6 biological replicates for each strain, each replicate is the mean of two technical replicates. *** p-value <0.0005, Mann-Whitney test, comparing the indicated mutant to WT. **C.** 96-well plate biofilm assay after 48h growth in BHIS. Mean of WT is adjusted to 100 %. Min-max boxplot of 6-9 biological replicates for each strain, each replicate is the mean of two technical replicates. *** p-value <0.0005, Mann-Whitney test, comparing the indicated mutant to WT. The images shown under each boxplot correspond to representative CV-stained microtiter wells after resuspension of the biofilm.

### *B. thetaiotaomicron* biofilm formation is modulated by capsule cross-regulation

To uncover the mechanism of increased biofilm formation in a *ΔCPS4* strain, we performed a random transposon mutagenesis in *ΔCPS4* and identified 6 mutants out of 4650 with reduced biofilm formation capacity compared to the parental *ΔCPS4* (Figure 2A). Five of these mutants corresponded to transposons inserted in the *BT1358-1357* region just upstream of the CPS4 operon (Figure 1A and Figure 2B, Supplementary Table S1). *BT1358* codes for an UpxY-like homolog and *BT1357* codes for a UpxZ–like homolog, two regulatory genes located at the beginning of most capsule synthesis operons in *B. thetaiotaomicron* and *B. fragilis* (18, 19). UpxY-like proteins positively regulate their cognate capsular operon by preventing premature transcription termination in the untranslated region, thus facilitating the otherwise abortive transcription of the downstream capsular genes (18). By contrast, UpxZ–like proteins are repressors of transcription of non-adjacent capsular systems (19). We first showed that deletion of *upxY^BT1358^* in *B. thetaiotaomicron ΔCPS4* did not impact biofilm formation, which is consistent with its role as a positive regulator of the expression of capsule 4 genes, all missing in the *ΔCPS4* mutant (Figure 2BC). We then hypothesized that transposon insertion in *upxY^BT1358^* (located upstream of *upxZ^BT1357^*) could have a polar effect on the expression of the repressor *upxZ^BT1357^* leading to the de-repression one or more of the 7 other *B. thetaiotaomicron* capsular polysaccharides. Indeed, in-frame deletion of *upxZ^BT1357^* or *upxY^BT1358^-upxZ^BT1357^* in a *ΔCPS4* background did not affect growth but led to loss of biofilm capacity (Figure 2C and Supplementary figure S1A). This phenotype could be complemented *in trans* by introducing *upxZB^T1357^* expressed from a constitutive promoter in the 5’ untranslated region of the tRNA-Ser chromosomal locus, either in *ΔupxZB^T1357^ΔCPS4* or *ΔupxY^BT1358^-upxZB^T1357^ΔCPS4 B. thetaiotaomicron* background (Figure 2C). To identify which capsules were repressed by *upxZB^T1357^*, we used qRT-PCR to monitor the expression of each capsular operon and we observed an increased transcription of capsule 2 (CPS2) in *B. thetaiotaomicron ΔupxZB^T1357^ΔCPS4* compared to *B. thetaiotaomicron ΔCPS4* single mutant (Supplementary Figure S2). Consistently, deletion of CPS2 operon in *B. thetaiotaomicron ΔupxZB^T1357^ΔCPS4* background restored biofilm formation capacity (Figure 2D) Thus, expression of either CPS4, or CPS2 in absence of CPS4, hinders biofilm formation.

**Figure 2.**
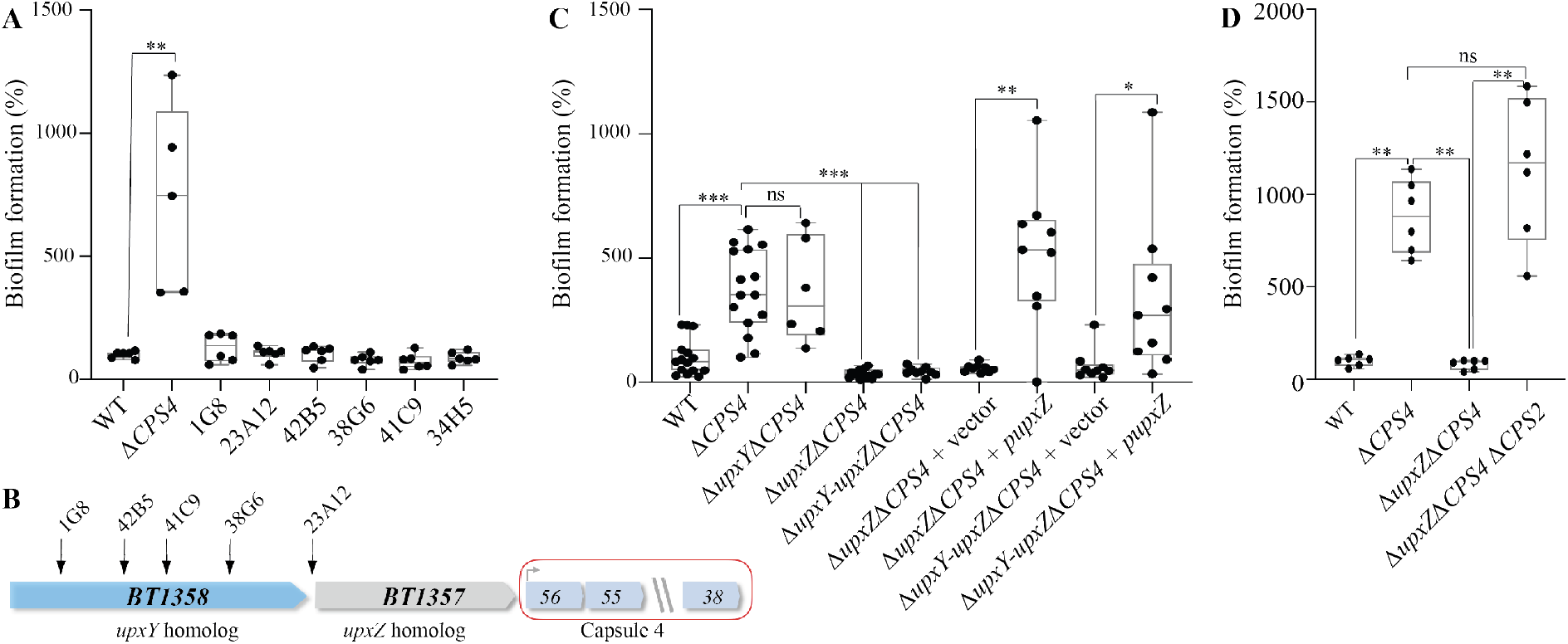
Capsule cross-regulation modulates biofilm formation in *B. thetaiotaomicron*. **A.** 96-well plate crystal violet biofilm assay after 48h growth in BHIS. **B.** Organization of *B. thetaiotaomicron* capsular operon 4 (CPS4) with identified transposon insertion points in the first two genes of the operon (*BT1358* and *BT1357*), coding regulators of capsular biosynthesis. **C.** 96-well plate crystal violet biofilm assay after 48h growth in BHIS. Mean of WT is adjusted to 100 %. **D.** 96-well plate crystal violet biofilm assay after 48h growth in BHIS. **A,C,D**: Mean of WT is adjusted to 100 %. Min-max boxplot of 6-9 biological replicates for each strain, each replicate is the mean of two technical replicates. ** p-value <0.005, Mann-Whitney test. **C.** and **D.** *upxY* strands for *upxY^BT1358^* and *upxZ* stands for *upxZ^BT1357^*.

### Expression of capsule 8 and lack of any capsules both induce biofilm formation

To assess the contribution of all capsules, besides inhibition by CPS4 or CPS2, to *B. thetaiotaomicron* biofilm formation, we used a recently described set of strains only expressing one of the eight *B. thetaiotaomicron* capsular types (20). We observed that derivative strains expressing only capsule 1, 2, 3, 4, 5 or 6 formed as little biofilm as wildtype (WT) *B. thetaiotaomicron* VPI5482. Interestingly strains only expressing CPS7 or CPS8 formed over 35 times more biofilm than the WT strain (Figure 3A). However, all CPS7-only bacteria seemed to be acapsulated, which is consistent with previous observations suggesting that capsule 7 may not be expressed in tested laboratory conditions (Supplementary Figure S3AB) (20). Indeed, similarly to a CPS7-only strain, a strain deleted for all 8 capsule operons (*ΔCPS1-8*) formed 40 times more biofilm than WT (Figure 3A) and showed strong aggregation phenotype in overnight cultures (Supplementary figure S3B). By contrast, India ink staining confirmed the presence of a capsule in biofilm-forming (but not aggregating) CPS8-only bacteria, suggesting that capsule 8 could have intrinsic adhesive properties (Supplementary Figure S3AB). These results showed that acapsulated cells have a strong adhesion capacity and that, except for CPS8 and potentially CPS7, the expression of all capsules hinders *B. thetaiotaomicron* biofilm formation.

**Figure 3.**
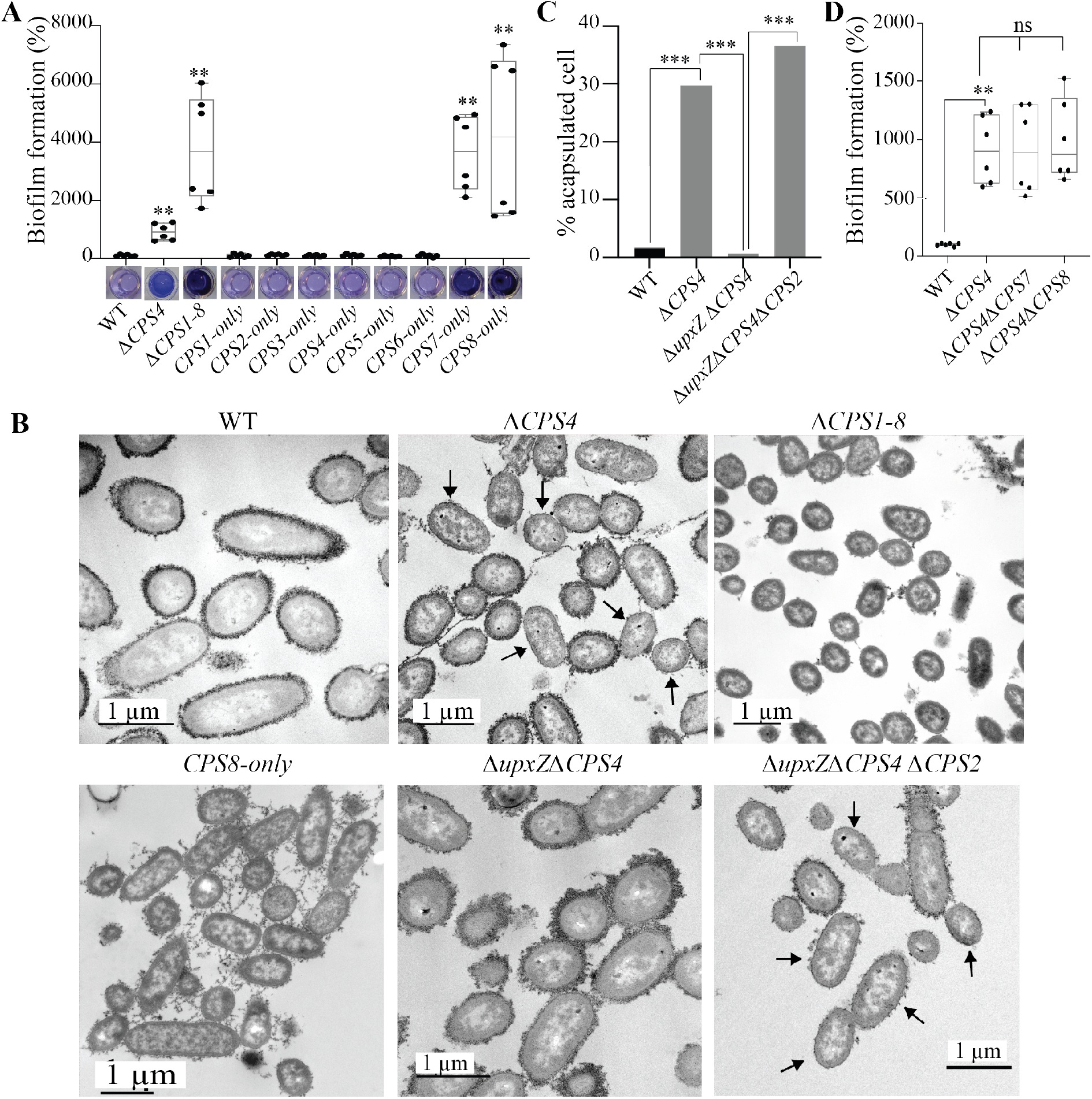
Capsule expression in *B. thetaiotaomicron* is heterogenous and has consequences on biofilm formation. **A.** and **D.** 96-well plate biofilm assay after 48h growth in BHIS. Mean of WT is adjusted to 100 %. Min-max boxplot of 6 biological replicates for each strain, each replicate is the mean of two technical replicates. ** p-value <0.005, Mann-Whitney test, comparing the indicated mutant to WT. The pictures shown under boxplot **A.** correspond to representative CV-stained microtiter wells after resuspension of the biofilm. **B.** Transmission electron microscopy (TEM) images of overnight cultures fixed with ferritin. Arrows indicate some example of acapsulated cells. **C.** Percentage of acapsulated cells of indicated strain counted on TEM pictures. For each strain at least 100 cells were counted. *** p-value<0.0005, prop.test (R). **B.** and **C.** *upxY*strands for *upxY^BT1358^* and *upxZ* stands for *upxZ^BT1357^*.

### Deletion of capsule 4 leads to a heterogeneously capsulated bacterial population

To determine whether lack of capsule or expression of the biofilm-promoting capsule 8 was responsible for the observed increased biofilm formation in *ΔCPS4* strain, we used Transmission Electron Microscopy (TEM) and showed that whereas WT *B. thetaiotaomicron* bacteria were almost all capsulated (>98%), ca. 30% of *ΔCPS4* cells lacked a visible capsule (Figure 3BC). Considering that *ΔCPS1-8* formed 4 times more biofilm than ΔCPS4, this suggested a correlation between increased frequency of non-capsulated cells in the population and the increased ability to form biofilms (Figure 3A and C). To determine whether capsulated cells in *ΔCPS4* population contributed to adhesion, we deleted, in the *ΔCPS4* background, either *CPS8*, the only biofilm-promoting capsule of *B. thetaiotaomicron*, or *CPS7*, for which we could not ascertain the biofilm formation potential using single CPS expressing strain. Both *ΔCPS4ΔCPS7* and *ΔCPS4ΔCPS8* mutants had similar biofilm capacity compared to a *ΔCPS4* mutant, showing that neither capsule 7 nor 8 contribute to biofilm formation in absence of capsule 4 (Figure 3D). Moreover, TEM imaging showed that the non-biofilm forming *ΔupxZ^BT1357^ΔCPS4* double mutant was entirely capsulated (due to induction of CPS2, Supplementary figure S2), supporting a correlation between increased biofilm formation (Figure 3A) and presence of a subpopulation of acapsulated cells in the *ΔCPS4* strain (Figure 3B and C). Consistently, deletion of *CPS2* in the *ΔupxZ^BT1357^ΔCPS4* background led to the apparition of 37% of acapsulated bacteria in a *ΔupxZ^BT1357^ΔCPS4ΔCPS2* population (Figure 3B and C) and restored biofilm formation (Figure 2D).

### Identification of BT2934 as a new *B. thetaiotaomicron* inhibitor of capsule expression

In addition to mutation in *ΔupxZ^BT1357^* capsule repressor, we also identified an additional biofilm-deficient *ΔCPS4* transposon mutant (34H5) with an insertion in *BT2934* (Figure 3A and 4A). *BT2934-2947* region corresponds to a *B. thetaiotaomicron* protein glycosylation locus (21, 22), in which *BT2934* encodes a homolog of the transmembrane oligosaccharide flippase Wzx (Figure 4A). We deleted *BT2934* and the 4 putative glycosyl transferases genes *BT2935-2938* located in the same operon and confirmed the role of *BT2934-2938* in protein glycosylation, as several bands disappeared from a protein glycosylation profile in *ΔCPS4ΔBT2934-2938* and 34H5 mutants compared to *ΔCPS4* (Supplementary figure S4). The double mutant *ΔCPS4 ΔBT2934-2938* had no growth defect and displayed a 2-fold decrease in biofilm formation compared to *ΔCPS4* (Figure 4B and Supplementary figure S1B). However, it still formed more biofilm than the original 34H5 transposon mutant in *BT2934*. To determine the origin of this discrepancy, we only deleted *BT2935-2938* glycosyl transferases genes and did not observe reduced biofilm capacity compared to the *ΔCPS4* strain. Although we did not succeed in deleting *BT2934* alone, introduction of *pBT2934*, constitutively expressing *BT2934*, in 34H5 transposon mutant and *ΔCPS4ΔBT2934-38* restored biofilm formation, but still showed an altered protein glycosylation profile (Figure 4C and supplementary figure S4). These results suggested that *BT2934* impact on biofilm formation did not involve *BT2935-2938* and might not directly involve protein glycosylation. Finally, we showed that while *ΔCPS4* and *ΔCPS4ΔBT2935-2938* bacteria displayed similar level of acapsulated cells (30% and 28% respectively), *ΔCPS4ΔBT2934-2938* cells showed full, wildtype level of capsulation (Figure 4DE), reduced back down to over 50% of capsulated cells upon complementation by *pBT2934* (Figure 4DE). To identify whether *BT2934* directly inhibited capsule production, we overexpressed *BT2934* in each single CPS expressing strains. We hypothesized that overexpression of BT2934 in each of these strains could inhibit capsule expression and lead to acapsulation of the whole population, thus leading to aggregation in overnight cultures. However, none of the resulting strains aggregated, hinting that no capsules were directly inhibited by an overexpression of *BT2934*. Taken together, these results suggest that *BT2934* indirectly impacts capsule production in *B. thetaiotaomicron*, with consequences on its ability to form biofilm.

**Figure 4.**
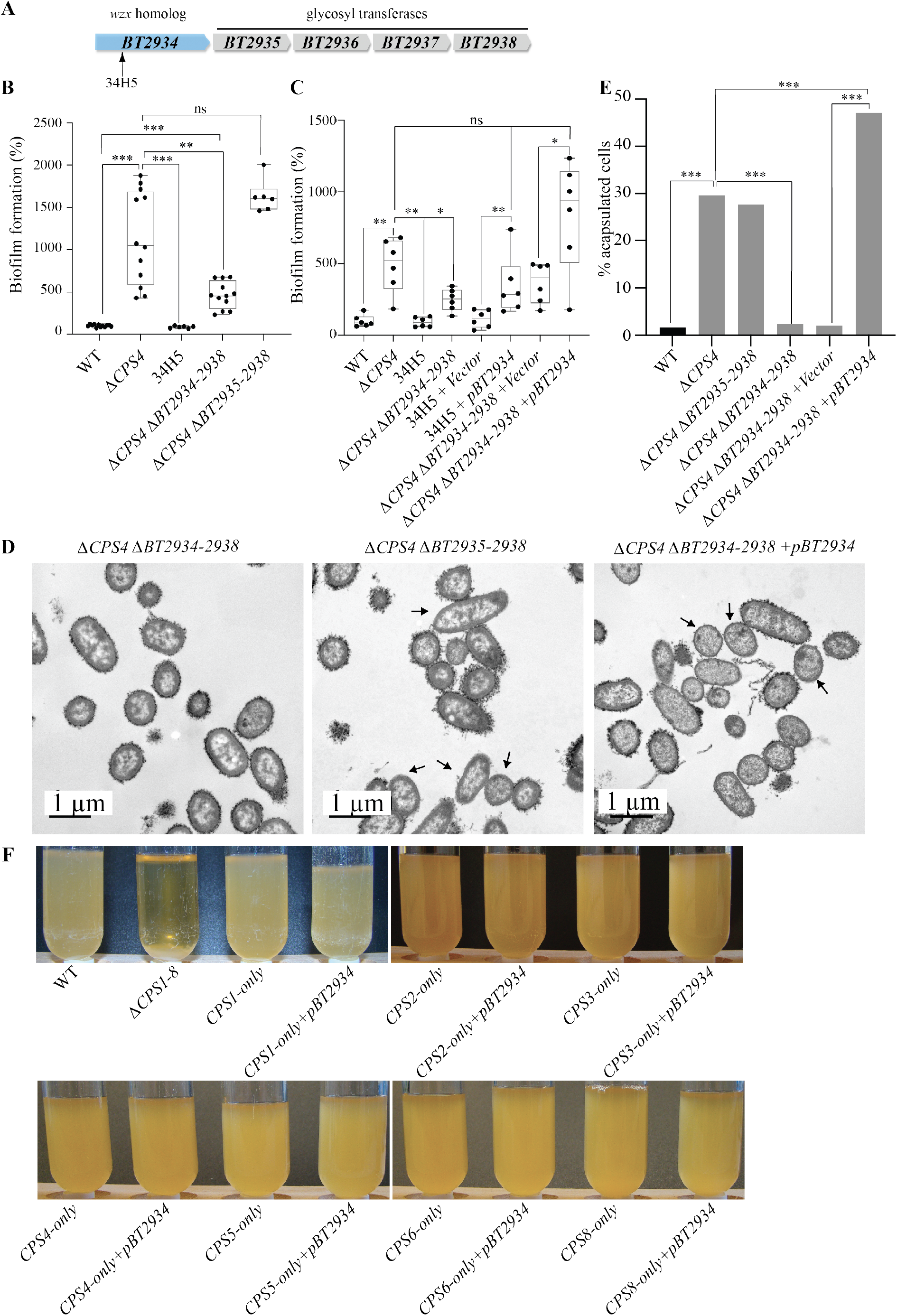
*BT2934* is a novel capsule inhibitor. **A.** Organization of *B. thetaiotaomicron* protein glycosylation *BT2934* locus with identified transposon insertion point. **B.** and **C.** 96-well plate crystal violet biofilm assay after 48h growth in BHIS. Mean of WT is adjusted to 100 %. Min-max boxplot of 6-12 biological replicates for each strain, each replicate is the mean of two technical replicates. *, p-value<0.05, ** p-value<0.005, *** p-value <0.0005, Mann-Whitney test. **D.** TEM images of *ΔCPS4ΔBT2934-2938, ΔCPS4ΔBT2935-2938* and *ΔCPS4ΔBT2934-2938+pBT2934* overnight cultures fixed with ferritin. Arrows indicate some acapsulated cells as an example. **E.** Percentage of acapsulated cells in overnight cultures counted on TEM pictures. For each strain at least 100 cells were counted. *** p-value<0.0005, prop.test (R). **F.** Overnight cultures of indicated strains in BHIS. Only ΔCPS1-8 showed aggregation.

### Biofilm-forming *CPS4* and *BT2934* mutants are outcompeted by the wildtype strain *in vivo*

*CPS4* and *BT2934* have previously been shown to be important for *in vivo* colonization in presence of a complex mix of *B. thetaiotaomicron* transposon mutants (23). To test whether unmasking *B. thetaiotaomicron* biofilm formation capacity could contribute to *in vivo* colonization, we used intragastric gavage to inoculate axenic mice with erythromycin-resistant WT-erm and tetracycline-resistant *ΔCPS4-tet or ΔBT2934-38-tet* in a 1:1 mix ratio and measured abundance of each strain in feces for 8 days using erythromycin and tetracycline resistance to discriminate between the strains. We first verified that *erm* and *tet* resistance markers did not impact *in vivo* colonization of WT-*erm* and WT-*tet* (Figure 5A). We then showed that both *ΔCPS4* and *ΔBT2934-38* were outcompeted by WT strain in two-strains cocolonization experiments (Figure 5BC), even though both *ΔCPS4* and *ΔBT2934-38* formed more biofilm than WT (Figure 1C and supplementary figure S6). When we tested colonization of the double mutant *ΔCPS4ΔBT2934-2938* against *ΔCPS4*, we found that they colonized mice similarly (Figure 5D), indicating that *BT2934* is only necessary for colonization in WT but not in *ΔCPS4* background. Taken together, these results showed that increased *in vitro* biofilm formation capacity is not predictive of *in vivo* colonization capacity.

**Figure 5.**
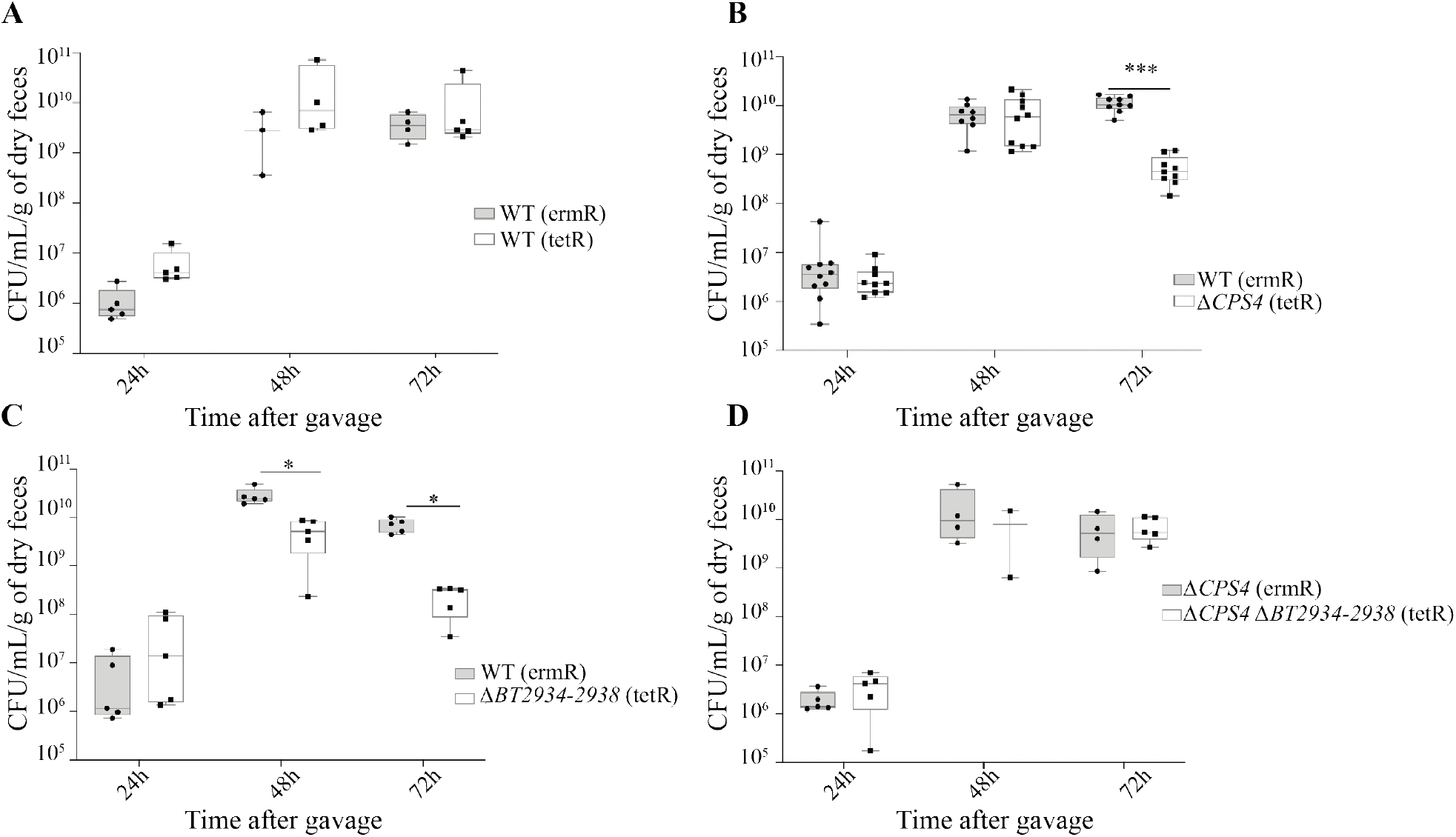
*BT2934* and *CPS4* contribute to *in vivo* colonization in axenic mice. Min-Max boxplot of CFU/mL/dry weight of feces, numbered from feces from 5-10 axenic mice after co-colonization with indicated strains. * pvalue<0.05, ** p-value<0.005, *** p-value<0.0005, Mann-Whitney test. **A.** WT (ermR) vs WT (tetR). **B.** WT (ermR) vs *ΔCPS4* (tetR). **C.** (ermR) vs *ΔBT2934-2938* (tetR). **D.** *ΔCPS4* (ermR) vs *ΔCPS4ΔBT2934-2938* (tetR).

## DISCUSSION

In contrast to oral *Bacteroidales*, intestinal *Bacteroidales* species possess numerous capsular polysaccharide loci that play important beneficial roles during gut colonization, ranging from protecting bacteria from stresses to mediating interactions with the host immune system (17, 20, 24–26). In this study we showed that deletion of one of *B. thetaiotaomicron* 8 capsular polysaccharides, CPS4, promotes biofilm formation *in vitro*, indicating that capsules mediate yet another important aspect of bacterial physiology.

Bacterial capsular polysaccharides are known to negatively affect biofilm formation by masking surface structures involved in adhesion in many bacteria (27–31). It was shown, for instance, that *Escherichia coli* capsular polysaccharides inhibit adhesion and autoaggregation by masking the short autotransporter adhesin antigen 43 as well as type III secretion system required for attachment in enteropathogenic *E. coli* (EPEC) (32, 33). The fact that CPS4 is the most expressed capsule in the tested laboratory conditions and *in vivo* (20) probably explains why it was the only capsule of *B. thetaiotaomicron* we identified by random transposon mutagenesis screening for increased biofilm formation.

In an adhering *ΔCPS4* strain, 30% of the bacteria are acapsulated, indicating that occurrence of only a subpopulation of acapsulated cells is enough to induce biofilm formation. In *Bacteroides fragilis*, acapsular cells were previously shown to aggregate (17, 34) and we also observed that a completely acapsular strain of *B. thetaiotaomicron* lacking all 8 capsules (*ΔCPS1-8*) displays a strong aggregation phenotype, suggesting cell-to-cell interactions driving biofilm formation in the absence of a capsule. However, due to the protective roles of *Bacteroides* capsules, acapsular strains are rapidly outcompeted by WT strain in axenic mice colonization (17, 34, 35). It is therefore unclear whether acapsular cells can be found *in vivo*, as studies following the expression of the 8 capsules of *B. thetaiotaomicron* by qRT-PCR would miss it, since there is no marker of acapsular cells. However, colonization of axenic mice with a mix composed of an acapsular mutant and 8 strains each expressing a single capsule showed that a low amount of acapsular cells was found to persist in the lumen of the small intestine of two out of five mice, potentially due to a decreased immune system pressure allowing the acapsular cells to survive (20).

*Bacteroides sp*. capsular loci are regulated by a complex transcriptional network, involving stochastic inversion of some capsule promoters (17, 36), transcriptional cross-regulation between capsular regulators UpxY and UpxZ (18, 19) and cross-talk between polysaccharide utilization loci and capsules through common sigma factors (37). It is also impacted by a range of environmental parameters such as diet, community composition and host physiology (20, 37, 38). In particular, expression of capsule 4 in mice has been shown to be increased *in vivo* compared to *in vitro* in a high fiber diet, but it is decreased in the suckling period compared to the weaned period (37, 38), and it is strongly impacted by the immune system (20). Moreover, a transcriptional analysis comparing planktonic cells with biofilms grown on chemostats for 8 days previously showed that CPS4 is downregulated in *B. thetaiotaomicron* biofilms (15).

Random transposon mutagenesis in *ΔCPS4* strain identified capsule regulation as the main parameter governing biofilm formation in our conditions. We show that *BT1357*, encoding the UpxZ homolog of *CPS4*, represses transcription of CPS2. As UpxZ proteins repress the transcription of non-adjacent capsular operon by interacting with the antiterminator UpxY proteins, necessary for the full transcription of their cognate capsules (18, 19), *BT1357* therefore most likely only interferes with *BT0462*, the UpxY homolog of *CPS2*. Whereas the complex interplay between UpxY and UpxZ homologs of *B. fragilis* was very well described, it is, to our knowledge, the first description of the precise inhibition pattern of a *B. thetaiotaomicron* UpxZ homolog (18, 19).

In addition to *BT1357*, we have identified that deletion of *BT2934* impacted capsule production. *BT2934-2947* is the protein O-glycosylation locus of *B. thetaiotaomicron* (21, 22). This locus is composed of a *wzx* oligosaccharide flippase (*BT2934*) and glycosyl transferases and its homolog in *B. fragilis, BF4298-4306* locus, was shown to be required for both *in vivo* and *in vitro* fitness in *B. fragilis* (21, 22). Accordingly, *BT2934* was previously shown to be important in both *in vitro* and *in vivo* competition experiments between complex communities of *B. thetaiotaomicron* transposon mutants (23) and was recently described as a putative essential gene (39). Our results confirm both the role of the *ΔBT2934-2938* locus in protein glycosylation and the decreased colonization capacity of a *ΔBT2934-2938* mutant in axenic mice in competition with the WT strain. However, deletion of *BT2934-2938* in *ΔCPS4* background had no effect on the colonization capacity of this strain. Although we never succeeded to delete *BT2934* alone, deletion of *BT2934-2938* in WT and *ΔCPS4* background did not lead to any growth defect *in vitro*, suggesting that deleting *BT2935-2938* might somehow alleviate the fitness cost associated with loss of *BT2934*.

We showed that deletion of *BT2934* impacted capsule production independently of protein glycosylation, as complementation by *BT2934* is sufficient to restore ΔCPS4 biofilm formation phenotype, but not the lack of protein glycosylation. The mechanism by which *BT2934* impacts capsule production remains to be elucidated. Because overexpression of *BT2934* in each single CPS expressing strains did not lead to general acapsulation, we hypothesize that *BT2934* does not directly inhibit capsule production. *BT2934* catalyzes the flipping of an oligosaccharide bound to an undecaprenyl-phosphate molecule across the membrane. As oligosaccharide flipping is also required for lipopolysaccharide and capsular synthesis, we speculate that these three processes might compete for undecaprenyl-phosphate or sugar moieties availability. Thus, limiting protein glycosylation by removing *BT2934* could favor the production of some capsules.

While our random transposition mutagenesis in *ΔCPS4* was not saturating, it is surprising that all identified biofilm-deficient mutants corresponded to insertions affecting capsule production rather than a putative adhesion factors unmasked in acapsulated bacteria. This could be indicative of the role played by purely electrostatic interactions between acapsulated bacteria or mediated by multiple and potentially redundant adhesive surface structures

We show that expression of all capsular polysaccharide of *B. thetaiotaomicron* hindered biofilm formation, except for CPS8 that rather promoted biofilm formation. Consistently, CPS8 expression was shown to be up-regulated in 8-day chemostat-grown biofilms (15), while capsules 1, 3, 4 and 6 were down-regulated. CPS8 might either be an adhesive capsule or a loose capsule that does not mask adhesion factors. However, if CPS8 did not mask adhesion factors we would expect CPS8-only strain to adhere like *ΔCPS1-8*, but CPS8-only formed less biofilm than *ΔCPS1-8* and it did not aggregate overnight. This suggests that capsule 8 could be a capsule providing adhesion capacity on its own. Interestingly, CPS8 is the only capsular locus of *B. thetaiotaomicron* containing homologs of FimA, the major component of type V pilus (40). Type V pili are widely found in *Bacteroidetes* and they were shown to mediate adhesion in *Porphyromonas gingivalis* (41, 42). Moreover, we previously showed that another homolog of FimA, BT3147, mediated biofilm formation in *B. thetaiotaomicron* upon truncation of the last 9 amino acids (13). CPS8 is expressed to low levels in axenic mice mono-colonized with *B. thetaiotaomicron*, and to slightly higher levels in mice colonized with complex communities, suggesting it might confer an advantage to *B. thetaiotaomicron* when competing with other bacteria for colonization. Whereas a strain expressing only CPS8 is rapidly outcompeted by the WT in *in vivo* competition experiment, some population of CPS8-only bacteria can be found in the lumen of the small intestine in some mice, reminiscent of the acapsular strain localization (20).

In this study, we found no evidence that higher *in vitro* adhesion would lead to better colonization of axenic mice. However, we assessed abundance of each strain by enumerating bacteria in the feces of mice, even though feces composition only partially recapitulates gut microbiota composition (43). In particular, we can imagine that cells with higher adhesion would not be shed in the feces as much as cells with low adhesion, mimicking a colonization defect. Moreover, besides biofilm formation, *CPS4* and *BT2934* participate in other significant processes *i.e*. interactions with the immune system and protein glycosylation respectively. Therefore, we cannot establish that the loss of *in vivo* fitness of *ΔCPS4* and *ΔBT2934-38* compared to WT is due to biofilm formation defect or to the loss of other functions impacted by the deletion of these genes.

In this study, we have shown that capsule regulation is a major determinant of biofilm formation and that competition between protein glycosylation and capsule production could constitute another layer of an already very complex capsule regulatory system. Further investigation of the mechanisms of biofilm formation in the gut commensal *B. thetaiotaomicron* will allow us to address the physiological adaptations of these bacteria within an anaerobic biofilm.

## MATERIALS AND METHODS

### Bacterial strains and growth conditions

Bacterial strains used in this study are listed in Table S2. *B. thetaiotaomicron* was grown in BHIS broth (44) supplemented with erythromycin 15 μg/ml (erm), tetracycline 2.5 μg/ml (tet), gentamycin 200 μg/ml (genta) or 5’-fluoro-2’-deoxyruidin 200 μg/ml (FdUR) when required and incubated at 37°C in anaerobic conditions using jars with anaerobic atmosphere generators (GENbag anaero, Biomerieux, ref. 45534) or in a C400M Ruskinn anaerobic-microaerophilic station. *Escherichia coli* S17λpir was grown in Miller’s Lysogeny Broth (LB) (Corning) supplemented with ampicillin (100 μg/ml) when required and incubated at 37°C with 180 rpm shaking. Cultures on solid media were done in BHIS with 1.5% agar and antibiotics were added when needed. Bacteria were always streaked from glycerol stock on BHIS-agar before being grown in liquid cultures. All media and chemicals were purchased from Sigma-Aldrich unless indicated otherwise. All experiments and genetic constructions of *B. thetaiotaomicron* were made in VPI *5482Δtdk* strain, which was developed for 2-step selection procedure of unmarked gene deletion by allelic exchange, as previously described (45). Therefore, the VPI *5482Δtdk* is referred to as wild type in this study.

### 96-well crystal violet biofilm formation assay

Overnight culture was diluted to OD_600_ = 0.05 in 100μL BHIS and inoculated in technical duplicates in polystyrene Greiner round-bottom 96-well plates. The wells at the border of the plates were filled with 200μL water to prevent evaporation. Incubation was done at 37°C in anaerobic conditions for 48h. The biofilm was fixed using 25μL of Bouin solution (picric acid 0.9%, formaldehyde 9% and acetic acid 5%, HT10132, Sigma-Aldrich) for 10min. Then the wells were washed once with water by immersion and flicking, and the biofilm was stained with 125uL 1% crystal violet (V5265, Sigma-Aldrich) for 10 minutes. Crystal violet solution was removed by flicking and biofilms were washed twice with water. Stained biofilms were resuspended in 1:4 acetone: ethanol mix and absorbance at 575nm was measured using TECAN infinite M200 PRO plate reader.

### Targeted mutagenesis

Deletion mutants were constructed using the previously described vector for allelic exchange in *B. thetaiotaomicron:* pExchange-*tdk* (45). A list of all the primers used in this study can be found in Table S3. Briefly, 1kb region upstream and downstream of the target sequence and pExchange-*tdk* were amplified by PCR using Phusion Flash High-Fidelity PCR Master Mix (Thermofischer Scientific, F548). All three fragments were ligated using Gibson assembly: the inserts and the plasmids were mixed with Gibson master mix 2x (100μL 5X ISO Buffer, 0.2 μL 10,000 U/mL T5 exonuclease (NEB #M0363S), 6.25 μL 2,000 U/mL Phusion HF polymerase (NEB #M0530S), 50 μL 40,000 U/mL Taq DNA ligase (NEB #M0208S), 87 μL dH2O for 24 reactions) and incubated at 50 °C for 35 min. The resulting mix was transformed in *E. coli* Sl7γpir that was used to deliver the vector to *B. thetaiotaomicron* by conjugation. Conjugation was carried out by mixing exponentially grown cultures (OD_600_=0.6) of the donor and the recipient strain in a 2:1 ratio. The mixture was spotted on BHIS-agar plates and incubated at 37°C in aerobic conditions overnight. The mix was then streaked on BHIS agar supplemented with antibiotic – for selection of *B. thetaiotaomicron* transconjugants that had undergone the first recombination event – and gentamicin to ensure exclusion of any *E. coli* growth. 8 of the resulting colonies were grown overnight in BHIS with no antibiotic to allow a second recombination event, and the culture was plated on BHIS-agar plates supplemented with FdUR to select for loss of plasmid. The resulting deletion mutants were confirmed by PCR and sequencing.

We used the pNBU2-bla-erm vector (46) for complementation, which inserts in the 5’ untranslated region of the tRNA-Ser, in which we previously cloned the constitutive promoter of *BT1311* encoding the sigma factor RpoD (13). We constructed a pNBU2-bla-tet vector by replacing the erythromycin resistance gene by a tetracycline resistance gene from the pExchange-tet plasmid using Gibson assembly (see above). Target genes were amplified by PCR using Phusion Flash High-Fidelity PCR Master Mix from start codon to stop codon and they were cloned after *BT1311* promoter by Gibson assembly. The Gibson mix was transformed in *E. coli* S17λpir and the resulting *E. coli* was used to transfer the plasmid to *B. thetaiotaomicron* by conjugation (see above).

### Transposon mutagenesis

pSAMbt, the previously published tool for random mariner-based transposon mutagenesis in *B. thetaiotaomicron* (23) was conjugated in *B. thetaiotaomicron* as described above. After streaking on BHIS-erm-genta agar plates, isolated colonies were resuspended in 100μL BHIS in 96-well plates, grown overnight and tested for biofilm formation as described above. The selected clones were then streaked on a fresh BHIS-erm-genta agar plate and 3 isolated colonies were tested for biofilm formation to ensure no mix of transposon mutants had occurred during preparation of the library. The genomic DNA of the validated clones was extracted using

DNeasy blood and tissue kit (Qiagen) and sent for whole genome sequencing at the Mutualized platform for Microbiology of Institut Pasteur.

### Electronic microscopy and numbering of acapsulated bacteria

Overnight cultures were adjusted to 1ml OD_600_=1.5. Cells were treated as described in Jacques and Foiry (Jacques and Foiry, 1989) for capsule observation: cultures were resuspended in glutaraldehyde 5% in 0.1M cacodylate buffer pH=7.2 and incubated at room temperature for 2h. Cells were then washed three times in 0.1M cacodylate buffer pH=7.2 and fixed 30min in 1mg/mL ferritin in 0.1M cacodylate buffer pH=7.2. Cells were washed one last time in 0.1M cacodylate buffer pH=7.2 and sent for transmission electronic microscopy at Electronic microscopy platform IBiSA of the University of Tours (https://microscopies.med.univ-tours.fr/). Acapsulated cells were counted by hand using Fiji cell counter plugin.

### India ink stain

5μL of overnight cultures were mixed with 3μL India ink directly on Superfrost plus glass microscopy slide (Thermofischer Scientific) and left to dry for 2min. The excess liquid was removed with paper towel after addition of the coverslip, and the cells were observed with a photonic microscope, 1000X.

### RNA extraction

Overnight cultures were mixed with RNA protect Bacteria Reagent (Qiagen) at a 1:2 volume ratio. The mix was incubated 5min at room temperature, then spun-down 10min 5000xg. The pellet was kept at −80°C. RNA was extracted from the pellet using FastRNA Pro Blue kit (MP). The pellet was resuspended in 1mL RNApro and mixed with lysing Matrix B. Cells were broken using FastPrep instrument at 40s, speed 6, twice at 4°C. The lysate was centrifugated for 10min at 4°C, 12000xg and the supernatant was collected and mixed with 300μL of chloroform. After 5min incubation at room temperature, the mix was centrifugated at 12000xg 4°C for 5min. The upper phase was transferred to a tube containing 500μL cold 100% ethanol and the nucleic acids were precipitated for 1h at −20°C. The tubes were centrifuged at 12000xg at 4°C for 15min and the pellet was washed in 500μL cold 75% ethanol. After centrifugation at 12000xg 4°C for 5min the ethanol was removed and the pellet was air-dried. 60μL of RNA-free water were added to the resuspend the nucleic acid and we treated it with TURBO DNAse from TURBO DNA free kit (Thermofischer Scientific, AM1907) for 1h30. Then the enzyme was inactivated using TURBO DNAse inactivator for 2min at room temperature and the extracted RNA was kept at −20°C.

### qRT-PCR

We performed reverse transcription using the First strand cDNA synthesis kit for RT-PCR (AMV) (Sigma-Aldrich) and the protocol described by the supplier. Briefly, 500μg RNA previously boiled 15min at 65°C were mixed with 2μL 10X reaction buffer, 4μL MgCl2 25mM, 2μL dNTP mix at 10mM each, 1μL 3’ primer 20μM, 1μL RNAse inhibitor 50U/μL, 0.8 AMV reverse transcriptase and water. The mix was incubated at 42°C for 1h30 and the enzyme was inactivated by heating to 99°C for 5min. qPCR was performed using SYBR green PCR master mix (Life technologies). cDNA was mixed with SYBR green master mix as described by supplier and with corresponding primers in technical duplicates in 384-well plates. qPCR reaction was performed using QuantStudio 6 Flex Real-Time PCR System (Thermofischer Scientific) and the “ΔΔCt method” program. We followed 16s rRNA and RpoB as housekeeping gene for normalization.

### Staining of glycosylated proteins

Overnight cultures were adjusted to 1mL OD_600_=1, spun-down and resuspended in 100μL 1X Laemmli-ß-mercaptoethanol lysis buffer (BioRad) and boiled 5min at 95°C. 10μL were run on Mini-PROTEAN TGX Stain-Free TM precast Gels (BioRad) in 1X TGX buffer for 40min at 170V. The gel was then stained using Pro-Q Emerald 300 staining for glycoproteins kit (Invitrogen) following the procedure described by the supplier.

### Co-colonization of axenic mice

Animal experiments were done at “Animalerie Axénique de MICALIS (ANAXEM)” platform (Microbiologie de l’Alimentation au Service de la Santé (MICALIS), Jouy-en-Josas, France) according to an official authorization n°3441-2016010614307552 delivered by the French ministry of Education nationale, enseignement supérieur et recherche. The protocol was approved by a local ethic committee on animal experimentation (committee n°45). All animals were housed in flexible-film isolators (Getinge-La Calhène, Vendôme, France) with controlled environment conditions (light/dark 12 h/12 h, temperature between 20 and 22°C, humidity between 45 to 55%). Mice were provided with sterile tap water and a gamma-irradiated standard diet (R03-40, S.A.F.E., Augy, France), *ad libitum*. Their bedding was composed of wood shavings and they were also given cellulose sheets as enrichment.

For each combination performed in separate isolators, groups of 5 male C3H/HeN germ free mice (6-11 weeks-old) were gavaged with 200μL bacterial suspensions containing 100 cells of each of the two strains we co-inoculated. One of the strains was conjugated with pNBU2-bla-erm plasmid, and the other with pNBU2-bla-tet plasmid so that they contained a different antibiotic resistance marker for later distinction. One of the combinations (WT (ermR) vs ΔCPS4 (tetR)) was performed twice (10 mice total). We plated the mix used for gavage onto BHIS agar plates with erythromycin or with tetracycline to check the initial ratio. When it was not 1:1, we corrected the measured abundance according to the ratio we had effectively used. At 24, 48 and 72 hours after inoculation, feces were collected, split in two tubes and weighed. One of the tubes was homogenized in 1 ml of BHIS, and serial dilutions were plated onto BHIS agar plates with erythromycin or with tetracycline. The abundance of each strain in the feces was measured by numbering the colony forming units growing on each type of plate using automated plater (easySpiral, Interscience, France) and counter (Scan500, Interscience, France). The other tube was dried in a speed vac concentrator (Savant, U.S.A.) and weighted. This allowed us to calculate the percentage of humidity of the feces we were using for each mouse and each condition and to infer the dry weight of feces used for CFU numbering. We divided the number of bacteria obtained by CFU counting by the dry weight of feces they were collected from.

### Growth curve

Overnight cultures were diluted to 0.05 OD_600_ in 200μL BHIS that had previously been incubated in anaerobic condition overnight to remove dissolved oxygen, in Greiner flat-bottom 96-well plates. A plastic adhesive film (adhesive sealing sheet, Thermo Scientific, AB0558) was added on top of the plate inside the anaerobic station, and the plates were then incubated in a TECAN Infinite M200 Pro spectrophotometer for 24 hours at 37°C. OD_600_ was measured every 30 minutes, after 900 seconds orbital shaking of 2 mm amplitude.

### Statistical analyses

Statistical analyses were performed using either R and Rstudio software or GraphPad Prism 8 for Mac OS (GraphPad software, Inc.). We used only non-parametric test. For *in vivo* experiments, 5-10 mice were used in 1 or 2 independent experiments. For all other experiments, at least 6 biological replicates in at least 2 independent experiment were used. A cut-off of p-value of 5 % was used for all tests. * p<0.05; ** p<0.05; *** p<0.005.

## ACKNOWLEDGEMENTS

We are grateful to Andy Goodman and Justin Sonnenburg for providing the genetic tools used in this study and to Eric Martens for generously providing the *B. thetaiotaomicron* capsule mutants and mono-expressing strains. We are also very thankful for Laurie Comstock comments on this manuscript. This work was supported by an Institut Pasteur grant and by the French government’s Investissement d’Avenir Program, Laboratoire d’Excellence “Integrative Biology of Emerging Infectious Diseases” (grant n°ANR-10-LABX-62-IBEID) and the *Fondation pour la Recherche Médicale* (grant no. DEQ20180339185). N. Béchon was supported by a MENESR (Ministère Français de l’Education Nationale, de l’Enseignement Supérieur et de la Recherche) fellowship. J. Mihajlovic was supported by the Pasteur Paris University International Doctoral Program and the *Fondation pour la Recherche Médicale* (grant no. FDT20160435523).

