## Supplementary material for "Capsular polysaccharides cross-regulation modulates *Bacteroides thetaiotaomicron* biofilm formation": SUPLEMENTAL FIGURES S1 TO S5

### SUPPLEMENTARY FIGURES

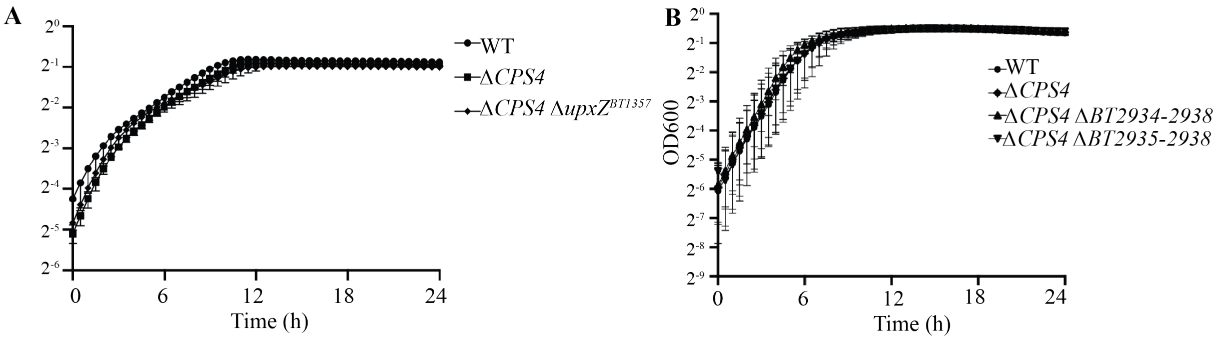


Supplementary Figure S1. **The mutants considered in this study were not affected for growth.** Planktonic growth of indicated mutants for 24h in BHIS in 96-well plates. OD600 was measured every 30min. Each dot is the median of 6 biological replicates. Error bars represent 95% confidence interval. **A.** Impact of *BT2934-2938* on growth. **B.** Δ*CPS1-8* transposon mutants.

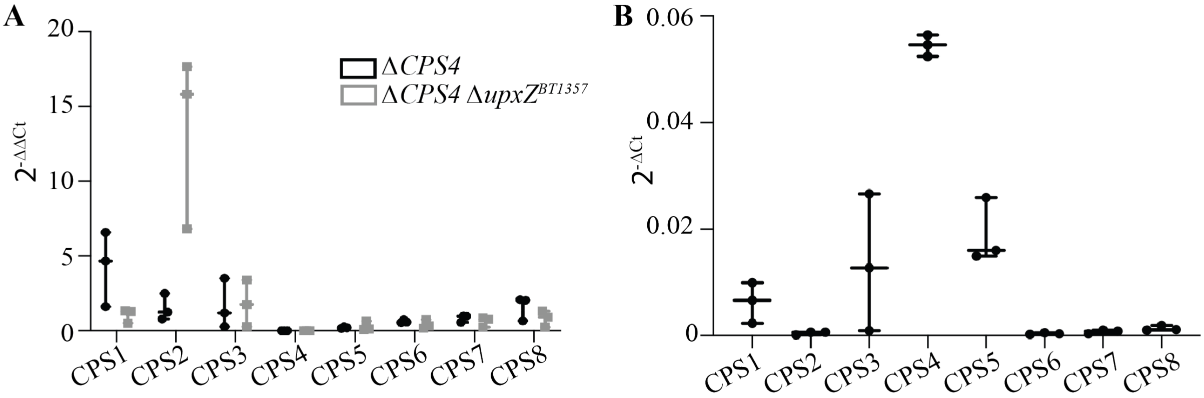


Supplementary Figure S2. ***BT1357* is a repressor of capsule 2 transcription.** qRT-PCR of 3 biological replicates of overnight planktonic cultures of WT, Δ*CPS4*, $\Delta CPS4 \Delta upxZ^{BT1357}$ following one gene of each capsular operon and 16sRNA and *rpoB* housekeeping genes. **A.** Expression levels of each capsules were controlled within each strain using the expression of 16s rRNA and *rpoB* housekeeping genes, and they were normalized between strains using the expression of WT as a reference. Y-axis represent fold-change of expression ($2^{-\Delta\Delta Ct}$) for each of the 8 capsules of *B. thetaiotaomicron.* B. Expression levels of each capsule normalized by 16sRNA and *rpoB* genes in WT. Y-axis represent fold-change of expression within one strain ($2^{-\Delta Ct}$) for each of the 8 capsules of *B. thetaiotaomicron.*

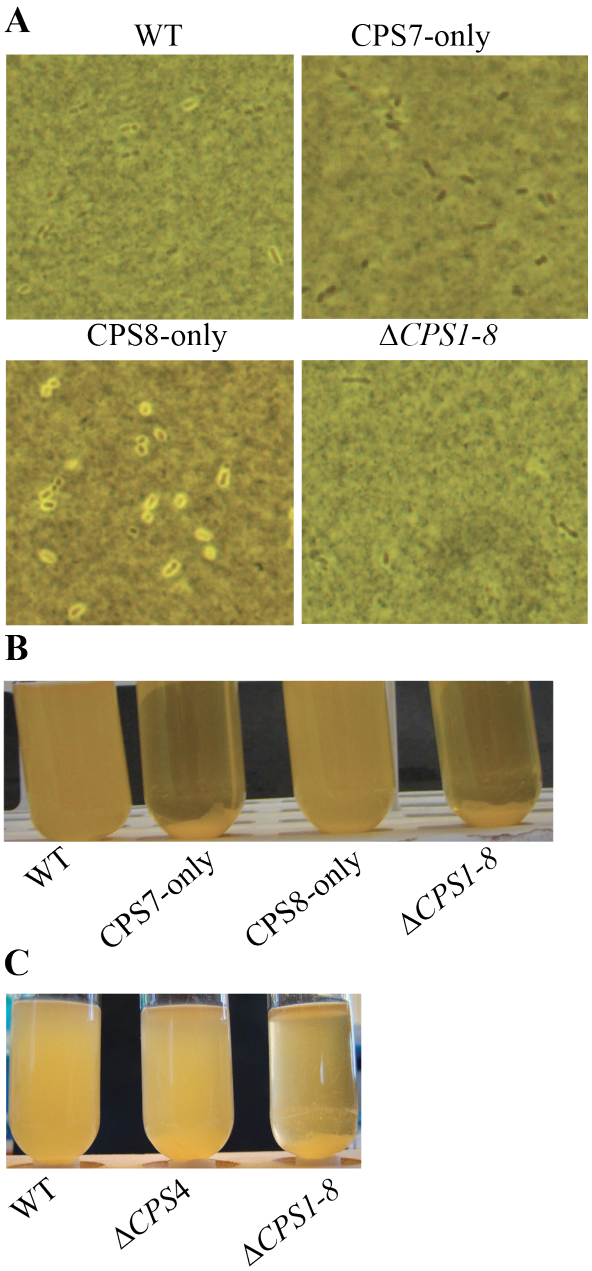


Supplementary figure S3. **Acapsulated *B. thetaiotaomicron* aggregate in BHIS. A.** India ink stained cultures observed under the phase contrast microscope, 1000X. **B.** and **C.** Aggregation in overnight cultures of *B. thetaiotaomicron* in BHIS.

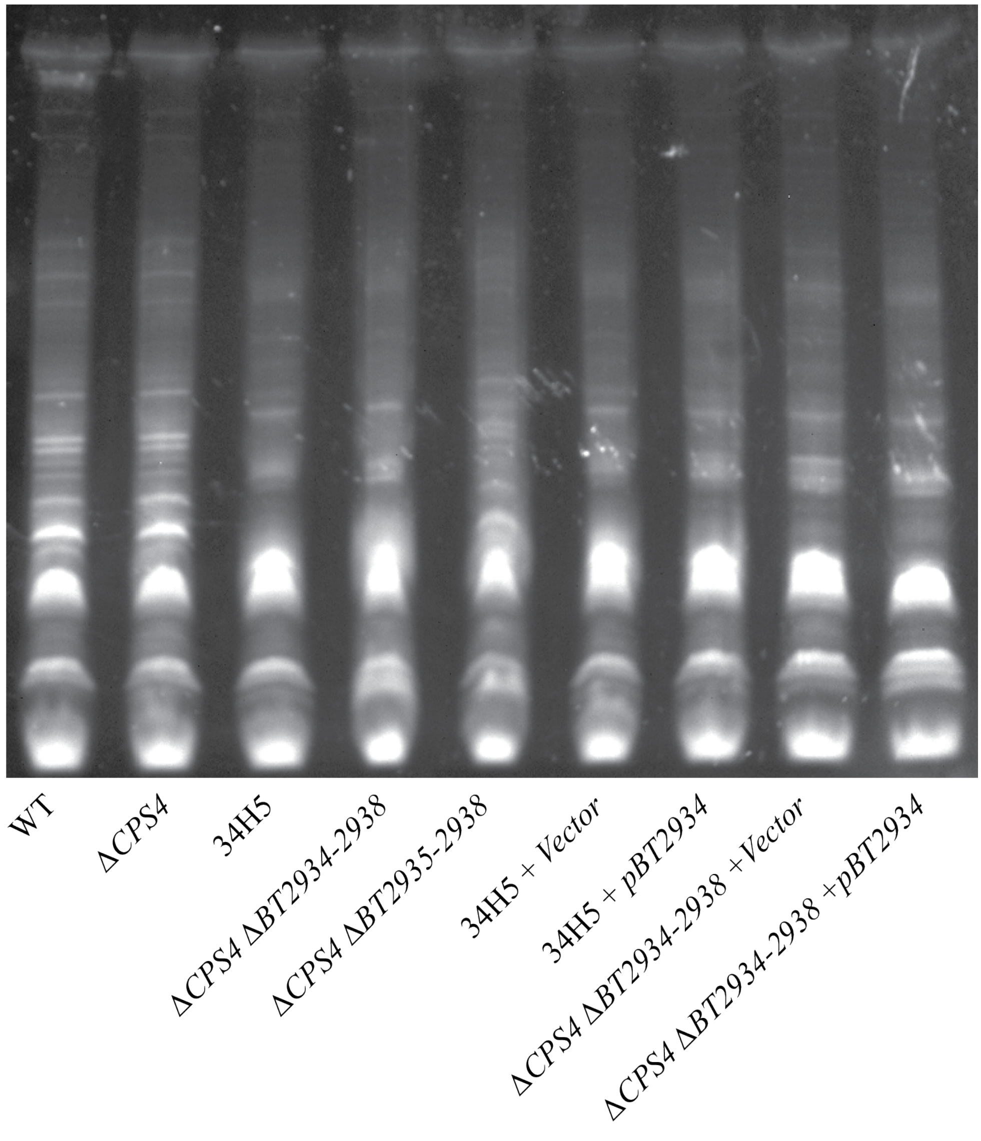


Supplementary Figure S4. ***BT2934-2938* is involved in protein glycosylation.** ProQ Emerald 300 staining of glycosylated proteins on whole cell extract of overnight cultures.

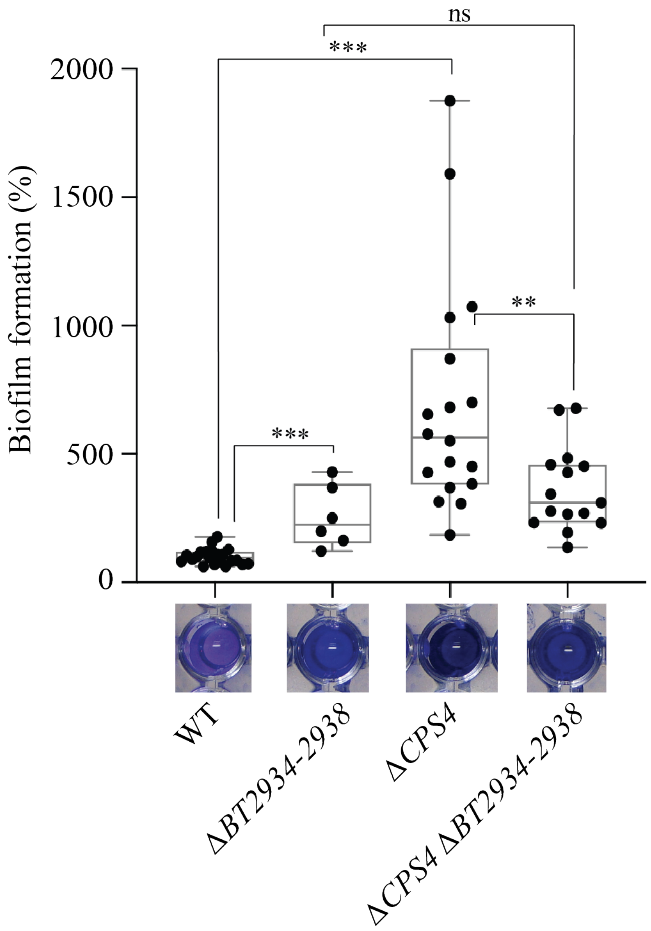


Supplementary figure S5. ***BT2934-2938* impacts biofilm formation**. **A.** 96-well plate crystal violet biofilm assay after 48h growth in BHIS. Mean of WT is adjusted to 100 %. Min-max boxplot of 6-18 biological replicates for each strain, each replicate is the mean of two technical replicates. * p-value<0.05, ** p-value <0.005, *** p-value <0.0005 Mann-Whitney test. The pictures shown under boxplot **C** correspond to representative CV-stained microtiter wells after resuspension of the biofilm.
