## Supplementary material for "Capsular polysaccharides cross-regulation modulates *Bacteroides thetaiotaomicron* biofilm formation": SUPLEMENTAL TABLES S1 TO S3

### SUPPLEMENTARY TABLES

Supplementary Table S1. **Transposon insertion targets.**

| **Name** | **Insertion** | **Gene annotation on NCBI** | **Function** | **Reference** |
|---|---|---|---|---|
| Transposon mutant in WT | | | | |
| 1C1 | *BT1356* | capsule polysaccharide export protein | CPS4 synthesis | Xu *et al.*, 2003 |
| 2C7 | *BT1342* | UDP-glucuronic acid epimerase | CPS4 synthesis | Xu *et al.*, 2003 |
| 3B12 | *BT1341* | UDP-glucose 6-dehydrogenase | CPS4 synthesis | Xu *et al.*, 2003 |
| 1G11 | *BT1353* | glycosyltransferase | CPS4 synthesis | Xu *et al.*, 2003 |
| 1F11 | *BT1352* | glycosyltransferase | CPS4 synthesis | Xu *et al.*, 2003 |
| Transposon mutant in Δ*CPS4* | | | | |
| 1G8 | *BT1358* | transcriptional regulator UpxY homolog | capsule regulation | Chatzidaki-Livanis *et al.*, 2010 |
| 23A12 | *BT1358 // BT1357* intergenic region | transcriptional regulator UpxY homolog // UpxZ homolog | capsule regulation | Chatzidaki-Livanis *et al.*, 2010 |
| 42B5 | *BT1358* | transcriptional regulator UpxY homolog | capsule regulation | Chatzidaki-Livanis *et al.*, 2010 |
| 38G6 | *BT1358* | transcriptional regulator UpxY homolog | capsule regulation | Chatzidaki-Livanis *et al.*, 2010 |
| 41C9 | *BT1358* | transcriptional regulator UpxY homolog | capsule regulation | Chatzidaki-Livanis *et al.*, 2010 |
| 34H5 | *BT2934* | wzx homolog, LPS biosynthesis transmembrane transport protein, Multidrug and toxin extrusion (MATE) domain | protein O-glycosylation | Fletcher *et al.*, 2009 |

1. Xu J, *et al.* (2003) A genomic view of the human-Bacteroides thetaiotaomicron symbiosis.
2. Chatzidaki-Livanis M, Weinacht KG, & Comstock LE (2010) Trans locus inhibitors limit concomitant polysaccharide synthesis in the human gut symbiont Bacteroides fragilis. *Proceedings of the National Academy of Sciences of the United States of America* 107(26):11976-11980.
3. Fletcher CM, Coyne MJ, Villa OF, Chatzidaki-Livanis M, & Comstock LE (2009) A general O-glycosylation system important to the physiology of a major human intestinal symbiont. *Cell* 137(2):321-331.
4. Martens EC, Roth R, Heuser JE, & Gordon JI (2009) Coordinate regulation of glycan degradation and polysaccharide capsule biosynthesis by a prominent human gut symbiont. *The Journal of biological chemistry* 284(27):18445-18457.

Supplementary Table S2. **List of strains and plasmids used in this study.**

| **Name in this paper** | **Genotype** | **Reference** |
|---|---|---|
| ***Bacteroides thetaiotaomicron*** | | |
| WT | VPI 5482 _*Δtdk* | Koropatkin *et al.*, 2008 |
| 1C1 | VPI 5482 _*Δtdk*- BT1356::Tn | This study |
| 2C7 | VPI 5482 _*Δtdk*- BT1342::Tn | This study |
| 3B12 | VPI 5482 _*Δtdk*- BT1341::Tn | This study |
| 1G11 | VPI 5482 _*Δtdk*- BT1353::Tn | This study |
| 1F11 | VPI 5482 _*Δtdk*- BT1352::Tn | This study |
| Δ*CPS4* | VPI 5482 _*Δtdk*_ΔBT1356-38 | This study |
| CPS1-only | VPI 5482 _*Δtdk*_Δ*cps2-8 cps1-lock* | Porter et al. 2017 |
| CPS2-only | VPI 5482 _*Δtdk*_Δ*cps1*_Δ*cps3-8* | Porter et al. 2017 |
| CPS3-only | VPI 5482 _*Δtdk*_Δ*cps1-2*_Δ*cps4-8 cps3-lock* | Porter et al. 2017 |
| CPS4-only | VPI 5482 _*Δtdk*_Δ*cps1-3*_Δ*cps5-8* | Porter et al. 2017 |
| CPS5-only | VPI 5482 _*Δtdk*_Δ*cps1-4*_Δ*cps6-8 cps5-lock* | Porter et al. 2017 |
| CPS6-only | VPI 5482 _*Δtdk*_Δ*cps1-5*_Δ*cps7-8 cps6-lock* | Porter et al. 2017 |
| CPS7-only | VPI 5482 _*Δtdk*_Δ*cps1-6*_Δ*cps8* | Porter et al. 2017 |
| CPS8-only | VPI 5482 _*Δtdk*_Δ*cps1-7 cps8-lock* | Porter et al. 2017 |
| Δ*CPS1-8* | VPI 5482 _*Δtdk*_Δ*cps1-8* | Porter et al. 2017 |
| Δ*CPS4*Δ*CPS7* | VPI 5482 _*Δtdk*_Δ*BT1356-38*_Δ*BT2886-62* | This study |
| Δ*CPS4*Δ*CPS8* | VPI 5482 _*Δtdk*_Δ*BT1356-38*_Δ*BT0038-68* | This study |
| 1G8 | VPI 5482 _*Δtdk*_Δ*BT1356-38-BT1358*::Tn | This study |
| 23A12 | VPI 5482 _*Δtdk*_Δ*BT1356-38-BT1358-7* intergenic region::Tn | This study |
| 42B5 | VPI 5482 _*Δtdk*_Δ*BT1356-38-BT1358*::Tn | This study |
| 38G6 | VPI 5482 _*Δtdk*_Δ*BT1356-38-BT1358*::Tn | This study |
| 41C9 | VPI 5482 _*Δtdk*_Δ*BT1356-38-BT1358*::Tn | This study |
| 34H5 | VPI 5482 _*Δtdk*_Δ*BT1356-38-BT2934*::Tn | This study |
| Δ*upxZ*$^{BT1357}$Δ*CPS4* | VPI 5482 _*Δtdk*_Δ*BT1356-38*_Δ*BT1357* | This study |
| *upxY*$^{BT1358}$Δ*CPS4* | VPI 5482 _*Δtdk*_Δ*BT1356-38*_Δ*BT1358* | This study |
| Δ*upxY*$^{BT1358}$-Δ*upxZ*$^{BT1357}$Δ*CPS4* | VPI 5482 _*Δtdk*_Δ*BT1356-38*_Δ*BT1358-7* | This study |
| Δ*upxZ*$^{BT1357}$Δ*CPS4*Δ*CPS2* | VPI 5482 _*Δtdk*_Δ*BT1356-38*_Δ*BT1357*_Δ*BT0463-82* | This study |
| Δ*upxY*$^{BT1358}$-Δ*upxZ*$^{BT1357}$Δ*CPS4* + *vector* | VPI 5482 _*Δtdk*_Δ*BT1356-38*_Δ*BT1358-7* + pNBU2-bla-erm-p1311 | This study |
| Δ*upxY*$^{BT1358}$-Δ*upxZ*$^{BT1357}$Δ*CPS4* + *pupxZ*$^{BT1357}$ | VPI 5482 _*Δtdk*_Δ*BT1356-38*_Δ*BT1358-7* + pNBU2-bla-erm-p1311-*BT1357* | This study |

| | | |
|---|---|---|
| Δ*upxZ*$^{BT1357}$Δ*CPS4* + *vector* | VPI 5482 *_Δtdk_ΔBT1356-38_ΔBT1357* + pNBU2-bla-erm-p1311 | This study |
| Δ*upxZ*$^{BT1357}$Δ*CPS4* + *pupxZ*$^{BT1357}$ | VPI 5482 *_Δtdk_ΔBT1356-38_ΔBT1357* + pNBU2-bla-erm-p1311-*BT1357* | This study |
| Δ*CPS4*Δ*BT2934-2938* | VPI 5482 *_Δtdk_ΔBT1356-38_ΔBT2934-8* | This study |
| Δ*CPS4*Δ*BT2935-2938* | VPI 5482 *_Δtdk_ΔBT1356-38_ΔBT2935-8* | This study |
| Δ*CPS4*Δ*BT2934-2938* + *Vector* | VPI 5482 *_Δtdk_ΔBT1356-38_ΔBT2934-8*+ pNBU2-bla-tet-p1311 | This study |
| Δ*CPS4*Δ*BT2934-2938* + *pBT2934* | VPI 5482 *_Δtdk_ΔBT1356-38_ΔBT2934-8*+ pNBU2-bla-tet-p1311-*BT2934* | This study |
| 34H5 + *Vector* | VPI 5482 *_Δtdk_ΔBT1356-38_BT2934*::Tn+ pNBU2-bla-tet-p1311 | This study |
| 34H5 + *pBT2934* | VPI 5482 *_Δtdk_ΔBT1356-38_BT2934*::Tn+ pNBU2-bla-tet-p1311-*BT2934* | This study |
| WT-ery | VPI 5482 *_Δtdk*+ pNBU2-bla-ery-p1311 | This study |
| WT-tet | VPI 5482 *_Δtdk*+ pNBU2-bla-tet-p1311 | This study |
| Δ*CPS4-ery* | VPI 5482 *_Δtdk_ΔBT1356-38*+ pNBU2-bla-ery-p1311 | This study |
| Δ*CPS4-tet* | VPI 5482 *_Δtdk_ΔBT1356-38*+ pNBU2-bla-tet-p1311 | This study |
| Δ*BT2934-2938* | VPI 5482 *_Δtdk_ΔBT2934-8* | This study |
| Δ*BT2934-2938 tet* | VPI 5482 *_Δtdk_ΔBT2934-8* + pNBU2-bla-tet-p1311 | This study |
| Δ*CPS4*Δ*BT2934-2938 tet* | VPI 5482 *_Δtdk_ΔBT1356-38_ΔBT2934-8*+ pNBU2-bla-tet-p1311 | This study |
| *CPS1-only+pBT2934* | VPI 5482 *_Δtdk_Δcps2-8 cps1-lock*+ pNBU2-bla-tet-p1311-*BT2934* | This study |
| *CPS2-only+pBT2934* | VPI 5482 *_Δtdk_Δcps1_Δcps3-8*+ pNBU2-bla-tet-p1311-*BT2934* | This study |
| *CPS3-only+pBT2934* | VPI 5482 *_Δtdk_Δcps1-2_Δcps4-8 cps3-lock*+ pNBU2-bla-tet-p1311-*BT2934* | This study |
| *CPS4-only+pBT2934* | VPI 5482 *_Δtdk_Δcps1-3_Δcps5-8*+ pNBU2-bla-tet-p1311-*BT2934* | This study |
| *CPS5-only+pBT2934* | VPI 5482 *_Δtdk_Δcps1-4_Δcps6-8 cps5-lock*+ pNBU2-bla-tet-p1311-*BT2934* | This study |
| *CPS6-only+pBT2934* | VPI 5482 *_Δtdk_Δcps1-5_Δcps7-8 cps6-lock*+ pNBU2-bla-tet-p1311-*BT2934* | This study |
| *CPS8-only+pBT2934* | VPI 5482 *_Δtdk_Δcps1-7 cps8-lock*+ pNBU2-bla-tet-p1311-*BT2934* | This study |
| ***Escherichia coli*** | | |
| S17λpir_pSAM-bt | S17λpir_pSAM-bt | This study |
| S17λpir_pExchange-tdk | S17λpir_pExchange-tdk | This study |
| S17λpir_pExchange-BT1356-38 | S17λpir_pExchange-BT1356-38 | This study |
| S17λpir_pExchange-BT2886-62 | S17λpir_pExchange-BT2886-62 | This study |
| S17λpir_pExchange-BT0038-68 | S17λpir_pExchange-BT0038-68 | This study |
| S17λpir_pExchange-BT1357 | S17λpir_pExchange-BT1357 | This study |
| S17λpir_pExchange-BT1358 | S17λpir_pExchange-BT1358 | This study |
| S17λpir_pExchange-BT1358-7 | S17λpir_pExchange-BT1358-7 | This study |
| S17λpir_pExchange-BT0463-82 | S17λpir_pExchange-BT0463-82 | This study |

| | | |
|---|---|---|
| S17λpir_pExchange-BT2934-8 | S17λpir_pExchange-BT2934-8 | This study |
| S17λpir_pExchange-BT2935-8 | S17λpir_pExchange-BT2935-8 | This study |
| S17λpir_ pNBU2-bla-erm-p1311 | S17λpir_ pNBU2-bla-erm-p1311 | This study |
| S17λpir_pNBU2-bla-erm-p1311-BT1357 | S17λpir_pNBU2-bla-erm-p1311-BT1357 | This study |
| S17λpir_pNBU2-bla-tet-p1311 | S17λpir_pNBU2-bla-tet-p1311 | This study |
| S17λpir_pNBU2-bla-tet-p1311-BT2934 | S17λpir_pNBU2-bla-tet-p1311-BT2934 | This study |
| **Name** | **Description** | **Reference** |
| **Plasmids** | | |
| pSAM-bt | Random transposon mutagenesis | Goodman *et al.*, 2009 |
| pExchange-tdk | Site-directed unmarked gene deletion | Martens *et al.*, 2008 |
| pExchange-BT1356-38 | Site-directed unmarked gene deletion of BT1356-38 | This study |
| pExchange-BT2886-62 | Site-directed unmarked gene deletion of BT2882-62 | This study |
| pExchange-BT0038-68 | Site-directed unmarked gene deletion of BT0038-68 | This study |
| pExchange-BT1357 | Site-directed unmarked gene deletion of BT1357 | This study |
| pExchange-BT1358 | Site-directed unmarked gene deletion of BT1358 | This study |
| pExchange-BT1358-7 | Site-directed unmarked gene deletion of BT1358-7 | This study |
| pExchange-BT0463-82 | Site-directed unmarked gene deletion of BT0463-82 | This study |
| pExchange-BT2934-8 | Site-directed unmarked gene deletion of BT2934-8 | This study |
| pExchange-BT2935-8 | Site-directed unmarked gene deletion of BT2935-8 | This study |
| pNBU2-bla-erm | Complementation | Wang et al 2000 |
| Vector (erm) | Complementation | This study |
| pupxZ$^{BT1357}$ | Complementation with BT1357 | This study |
| Vector (tet) | Complementation | This study |
| pBT2934 | Complementation with BT2934 | This study |

Supplementary Table S**3. Primers used in this study**

| **Construct** | **Name** | **Sequence (5'-->3')** |
|---|---|---|
| pExchange | pEx-ch-F | TGGGAATTCCCCTCCACCGC |
| | pEx-ch-R | GGGGAGAGGACGGACAGAAGAT |
| | pExchangeR | CGTCGACTCGAATGTTATCTTC |
| | pExchangeF | TCTAGAGCGGCCGCCACC |
| pExchange-BT1356-1338 | 1356-5R | AGAACAAAAAAGTAGAATGCTAAAAAGTCGTTGTATTTTC |
| | 1356-5F | GCGGTCGACAGAATAACGAATAAGTGTGGGG |
| | 1356-3F | GAAAATACAACGACTTTTTAGCATTCTACTTTTTTGTTCTTTCTTAGAATTGAAATAAACG |
| | 1356-3R | CCCTCTAGAATTCGATCCTACGAATACGACC |
| pExchange-BT0038-0068 | cps8-intF | ACGGCATAGATGTAGCCAAAAG |
| | cps8-intR | GATCGCCCAACCTAGTGTC |
| | cps8-3F | TTGATAATTCTTTTAAACATGATGCATGGTCTATTACAACCTGTC |
| | cps8-3R | GCGGTGGCGGCCGCTCTAGACAAAGAAGCTTTCCTTCTGAC |
| | cps8-5R | GGTTGTAATAGACCATGCATCATGTTTAAAAGAATTATCAATAGAAACATG |
| | cps8-5F | AGATAACATTCGAGTCGACGCATAAATTAGTTTGAGCGACG |
| pExchange-BT2886-2862 | cps7-extF | GACTGTTCCGAACATTG |
| | cps7-extR | AGCCTTGCTCAAAAACTGG |
| | cps7-5R | GTCAGAGAGAAATGATAGCAAATAACTCCTCCAATTCTATCATTTAAAG |
| | cps7-5F | AGATAACATTCGAGTCGACGGCAGAGTGAACTTTATCCTC |
| | cps7-3F | ATAGAATTGGAGGAGTTATTTGCTATCATTTCTCTCTGACAGATG |
| | cps7-3R | GCGGTGGCGGCCGCTCTAGATCCTTAGTCCCTGTACCC |
| pExchange-BT1358 | 1358-5F | GCGGTGGCGGCCGCTCTAGACAGCAGCCTCATTCTTTAGTGC |
| | 1358-5R | TTTCACTTATATTTAAACCCCTATTTACCCCACACTTATTCGT |
| | 1358-3F | AATAAGTGTGGGGTAAATAGGGGTTTAAATATAAGTGAAACAAGCA |

| | | |
|---|---|---|
| | 1358-3R | AGATAACATTCGAGTCGACGGTTGCATGCACTCATCCG |
| pExchange-BT1357 | 1357-3R | AGATAACATTCGAGTCGACGTGATTGTAATCTTGATATGCTAGCAAAG |
| | 1357-3F | ATTAAGTGAATAAAAGGGGTCTTTTTATTTGTCAATGAAAATACAACGAC |
| | 1357-5R | TTCATTGACAAATAAAAAGACCCCTTTTATTCACTTAATGCTTG |
| | 1357-5F | GCGGTGGCGGCCGCTCTAGATGTCAAGTTCAGGTTTCAGGTC |
| pExchange-BT1358-1357 | 1357-58 3F | AATAAGTGTGGGGTAAATAGCTTTTTATTTGTCAATGAAAATACAACGAC |
| | 1357-58 5R | TTCATTGACAAATAAAAAGCTATTTACCCCACACTTATTCGT |
| pExchange-BT0463-0482 | cps2-3R | GCGGTGGCGGCCGCTCTAGAGATTGAAAGTGGCGGCAG |
| | cps2-3F | TGAGTACTAATAACAAATCATCCGGTTCTAAGAATAAACCTGAG |
| | cps2-5R | GGTTTATTCTTAGAACCGGATGATTTGTTATTAGTACTCAAGATTTGAG |
| | cps2-5F | AGATAACATTCGAGTCGACGTGAAAGCTGTCAGGAAGCTAAG |
| pExchange-BT2934-2938 | 2934-5F | AGATAACATTCGAGTCGACGGGAAAGACTTCCAGGCACG |
| | 2934-5R | GCATTTGGGGACTTCACCGGATTATTATAATCGGTGAAGGAGAGTG |
| | 2938-3F | CCTTCACCGATTATAATAATCCGGTGAAGTCCCCAAATGC |
| | 2938-3R | GCGGTGGCGGCCGCTCTAGATCTCTATGAATGTCCTGTTTTCGT |
| pExchange-BT2935-2938 | 2935-5F | AGATAACATTCGAGTCGACGGGATAGCTCCGCGAGCTC |
| | 2935-5R | GCATTTGGGGACTTCACCGGTCATGATATCTTTTTCTTTTTAATGAACTG |
| | 35-2938-3F | AAAAGAAAAAGATATCATGACCGGTGAAGTCCCCAAATGC |
| pNBU2-bla | pNBU-chR | GCCAATGCACAAATGCTGTTCC |
| | pNBU-chF | CAGGTGTATTCCCATCCGG |
| | pNBU-F | CGACGTCGACTAATTGCC |
| | pNBU-R | ATGTTAAAAACAGATTTGGAGTGC |

| | | |
|---|---|---|
| pNBU2-bla-tetR | pNBU-del-eryF | ATGTCATCAAAATAAAAACAATAGGCCACATGCAAC |
| | pNBU-del-eryR | ATTTATAATATTCATTATAACCTCTCCTTAATTTATTG |
| | tetQ-F | ATTAAGGAGAGGTTATAATGAATATTATAAATTTAGGAATTCTTGCTC |
| | tetQ-R | TTGCATGTGGCCTATTGTTTTTATTTTGATGACATTGATTTTTGG |
| | tetQ-NBU-chF | CGGGATGAACCATGAGTAC |
| | tetQ-NBU-chR | GTACCGAGGACGCGTAAAC |
| pNBU2-bla-erm-BT1357 | pNBU-1357F | TCCAAATCTGTTTTTAACATATGGTGAGTTTTTTACTACAAG |
| | pNBU-1357R | TAGGCAATTAGTCGACGTCGTTAGTTGGTTTCACACAGTTCC |
| pNBU2-bla-tet-p1311-BT2934 | NBU-2934-F | CACTCCAAATCTGTTTTTAACATATGAGTGAAGAACAGTCACTGAAAC |
| | NBU-2934-R | GATAGGCAATTAGTCGACGTCGTCATGATATCTTTTTCTTTTTAATGAACTG |
| qPCR | 16SQF | TCAGCTCGTGTTGTGAAATG |
| | 16SQR | GTAAGGGCCATGATGACTTG |
| | rpoB-Q-F | CAAATTTCACGCCCAAAGTT |
| | rpoB-Q-R | GTGCGTCAGGAAGATATG |
| | 0381-QF | CGCTTTATCATGTCGTTGGA |
| | 0381-QR | GTACAAGCCGGAGCTTTTTG |
| | 0463-QF | TCCGATCACAAAAGGAGTGA |
| | 0463-QR | ACGTTTAAAGCCCGGAAGAT |
| | 0602-QF | TTGGGTTACATCGGTCTTCC |
| | 0602-QR | TTTGAAGGTGGTGTGACCTG |
| | 1355-QR | CGCAGGTTCTATCACTGCAA |
| | 1355-QF | AGGAGCGATTGCAAAATGAC |
| | 1653-QF | TGTTCCATTGAAAGCTTCAGG |
| | 1653-QR | CTCCCCATCAATATCCAGCTT |
| | 1722-QR | TGCAAAGACTGGCTTTTCCT |
| | 1722-QF | TGAGCAGGAGCGTTTACAGA |
| | 2885-QR | TAGAGGAAGAGTCCCGTTGGT |
| | 2885-QF | GGCAACGCAAATGGATTACTA |
| | 0039-QF | GGCATTTGCTTGTTTTATGGA |
| | 0039-QR | AGGGCTGGTCATATCGTTTCT |
